# Cryo-EM of native membranes reveals an intimate connection between the Krebs cycle and respiration in mycobacteria

**DOI:** 10.1101/2024.10.09.617439

**Authors:** Justin M. Di Trani, Jiacheng Yu, Gautier M. Courbon, Ana Paula Lobez Rodriguez, Chen-Yi Cheung, Yingke Liang, Claire E. Coupland, Stephanie A. Bueler, Gregory M. Cook, Peter Brzezinski, John L. Rubinstein

## Abstract

Imaging of endogenous protein complexes in their native membranes can reveal protein-protein interactions that are lost upon detergent solubilization. To investigate interactions in the mycobacterial oxidative phosphorylation machinery, we prepared inverted membrane vesicles from *Mycobacterium smegmatis* and enriched for vesicles containing complexes of interest by affinity chromatography. Electron cryomicroscopy (cryo-EM) of these vesicles revealed that malate-quinone oxidoreductase (Mqo), an enzyme from the Krebs cycle, physically associates with the electron transport chain Complex III_2_IV_2_ (CIII_2_CIV_2_) supercomplex. Analysis of the Mqo:CIII_2_CIV_2_ interaction shows that CIII_2_CIV_2_ is necessary for malate-driven, but not NADH-driven, electron transport chain activity and oxygen consumption. Further, the association of Mqo with CIII_2_CIV_2_ enables electron transfer from malate to CIII_2_CIV_2_ with millisecond kinetics. Together, these findings indicate a connection between the Krebs cycle and respiration that directs electrons along a single branch of the mycobacterial electron transport chain.

## Introduction

Biological energy is extracted from nutrients through metabolic pathways including glycolysis, the tricarboxylic acid or Krebs cycle, and fatty acid oxidation. In most organisms, the Krebs cycle provides reduced nicotinamide adenine dinucleotide (NADH) and succinate to membrane-bound electron transport chain (ETC) complexes to drive production of a transmembrane proton motive force (pmf). The pmf, in turn, provides the energy for synthesis of adenosine triphosphate (ATP) from adenosine diphosphate (ADP) and inorganic phosphate (P_i_). NADH is oxidized by Complex I of the ETC, reducing ubiquinone to ubiquinol. Oxidation of succinate to fumarate is an essential reaction in the Krebs cycle but occurs through Complex II of the ETC, which also reduces ubiquinone to ubiquinol. Electrons from ubiquinol are then transferred sequentially to Complex III, cytochrome *c* (cyt. *c*), Complex IV, and then to oxygen, reducing it to water. Complexes I, III, and IV couple electron transfer to proton translocation across the membrane, maintaining the pmf that powers ATP synthesis.

Mycobacterial ETCs differ from the canonical mammalian mitochondrial ETC in a number of ways (reviewed in (Liang and Rubinstein, 2023)). First, mycobacterial ETCs rely on menaquinone (MQ) rather than ubiquinone. Further, unlike canonical ETCs, mycobacterial ETCs are highly branched. In most mycobacteria, such as the pathogen *Mycobacterium tuberculosis* and the fast-growing saprophyte *Mycobacterium smegmatis*, NADH:MQ oxidoreductase activity is catalyzed by both Complex I and one or more non-proton pumping type II NADH dehydrogenases (NDH-2s). Two different enzymes, Sdh1 and Sdh2, catalyze succinate:MQ oxidoreductase activity. Further, both *M. tuberculosis* and *M. smegmatis* possess a malate:quinone oxidoreductase (Mqo) that oxidizes malate to oxaloacetate, a critical step in the Krebs cycle, while reducing MQ to MQH_2_ (Harold et al., 2022). In *M. tuberculosis*, this Mqo is found in addition to malate dehydrogenase (MDH), which transfers electrons from malate to NAD^+^, while in *M. smegmatis* Mqo is the only malate oxidizing enzyme (Harold et al., 2022). Perhaps most strikingly, oxidation of MQH_2_ in mycobacteria is catalyzed by a supercomplex of Complexes III and IV (CIII_2_CIV_2_) with a bound cytochrome *cc* subunit taking the place of soluble cyt. *c*. Oxidation of MQH_2_ and reduction of oxygen to water can also be accomplished by a cytochrome *bd* complex, not found in canonical ETCs, that translocates fewer protons per electron transfer than CIII_2_CIV_2_ (Safarian et al., 2021).

Study of the structures and interactions of mycobacterial ETC enzymes, just like other membrane proteins, is typically complicated by the need to first extract the proteins from their native lipid bilayer with detergents (Gong et al., 2018, 2020; Liang et al., 2023; Safarian et al., 2021; Wang et al., 2021; Wiseman et al., 2018; X. Zhou et al., 2021). Recently, large membrane protein complexes in their native membrane were imaged and reconstructed to high resolution (Coupland et al., 2024; Wang et al., 2024; Zheng et al., 2024). However, these experiments were allowed by special cases where the protein of interest is found in high abundance in the membrane of a specific organelle, such as ETC complexes in mitochondria (Zheng et al., 2024) and vacuolar-type ATPases in synaptic vesicles (Coupland et al., 2024; Wang et al., 2024). Preparation of cell-derived vesicles following over-expression has also allowed structure determination for a protein in a non-native membrane (Tao et al., 2023).

In contrast, we prepared inverted membrane vesicles (IMVs) from *M. smegmatis* cytoplasmic membranes, which contain all the membrane proteins found in this organism. We then enriched for IMVs and membrane fragments containing ATP synthase or CIII_2_CIV_2_ with affinity tags on the proteins of interest. We demonstrate that it is possible to use electron cryomicroscopy (cryo-EM) to obtain structures of both mycobacterial ATP synthase and mycobacterial CIII_2_CIV_2_ in their native membrane, the latter to high resolution. Three-dimensional (3D) reconstruction shows that in the absence of detergents, Mqo binds CIII_2_CIV_2_. The presence of CIII_2_CIV_2_ is necessary for malate-driven acidification of IMVs and malate-driven oxygen consumption by IMVs. Further, stopped flow spectroscopy demonstrates that the association of Mqo with CIII_2_CIV_2_ enables electron transfer from malate to CIII_2_CIV_2_ with millisecond kinetics, while electron transfer from NADH to CIII_2_CIV_2_ via MQ occurs more slowly. Together, these findings reveal a surprising additional direct connection between the Krebs cycle and ETC in mycobacteria.

## Results

### Structure of mycobacterial ATP synthase from inverted membrane vesicles

To establish if we could locate and determine structures of mycobacterial membrane proteins in IMVs formed from native membranes, we first prepared vesicles from an *M. smegmatis* strain with 3×FLAG tags at the C termini of the ATP synthase β subunits encoded by the genomic DNA (Guo et al., 2021). IMVs were formed by homogenizing membranes in buffer, which produces vesicles capable of succinate-driven ATP synthesis (Courbon et al., 2023). Vesicles that include ATP synthase were enriched by FLAG affinity chromatography, purified further by size exclusion chromatography, and imaged by cryo-EM on graphene oxide-coated specimen grids (**Fig. 1A**, *upper*). ATP synthase complexes protruding from the vesicle membranes were selected and used to calculate two-dimensional (2D) class average images, which show the familiar structure of the enzyme (Courbon et al., 2023; Guo et al., 2021) (**Fig. 1A**, *lower*). Images that contributed to these 2D classes were then used to calculate a 3D map of ATP synthase at ∼9 Å resolution (**Fig. S1** and **S2**, **Table S1**). Resolution in the catalytic F_1_ region of the complex was improved to ∼8.1 Å by focused refinement. A composite map of the complex shows the known structure of the mycobacterial ATP synthase (Courbon et al., 2023; Guo et al., 2021), but the map does not reveal any unexpected features or interactions involving the enzyme (**Fig. 1B**).

**Figure 1.**
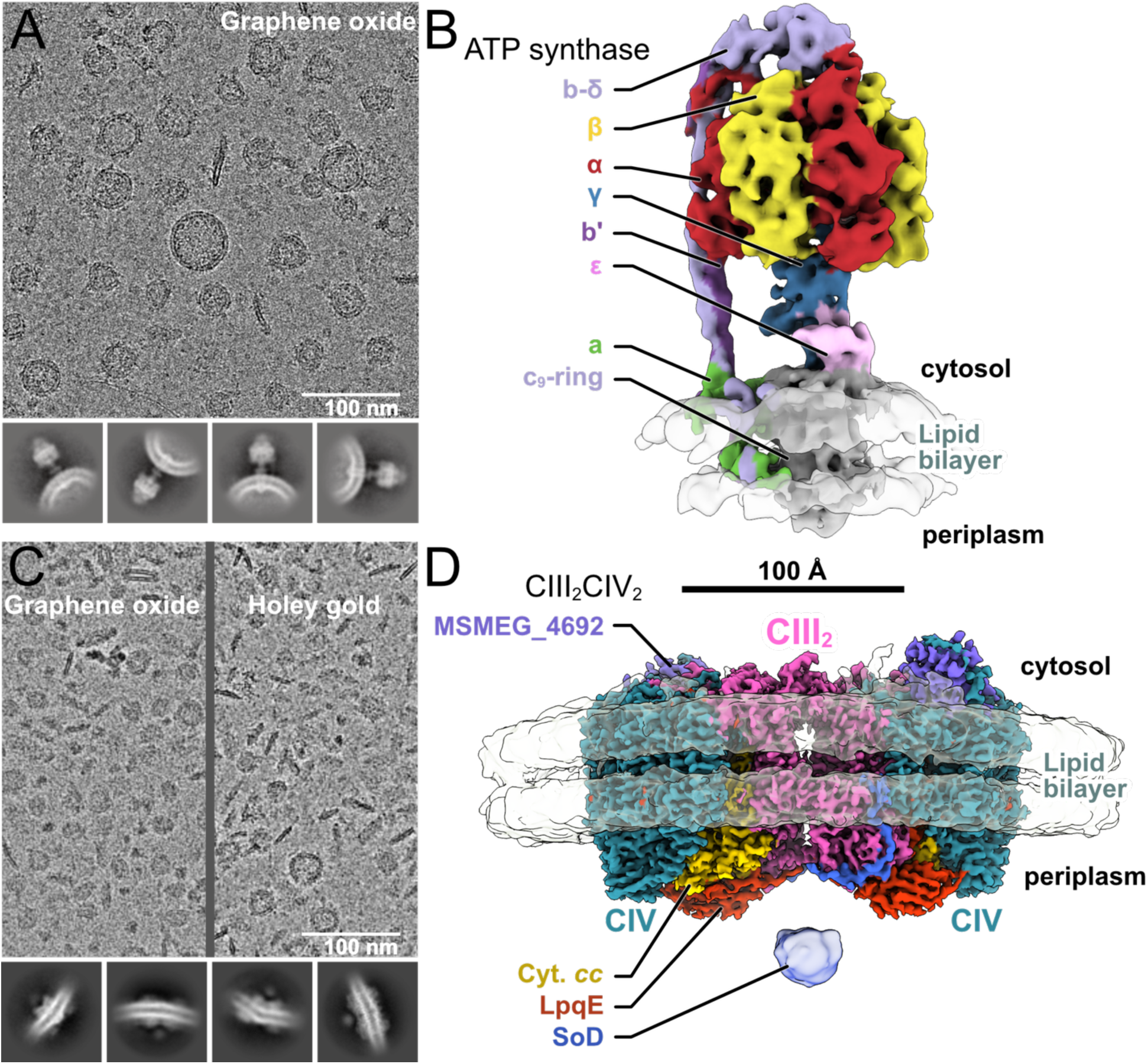
Structure determination of membrane proteins in native lipid bilayers. **A,** Cryo-EM micrograph and class average images from mycobacterial IMVs enriched for ATP synthase on graphene oxide-coated holey gold grids. **B,** Three-dimensional reconstruction of mycobacterial ATP synthase in its native membrane. **C,** Cryo-EM micrograph and class average images from mycobacterial IMVs and membrane fragments enriched for CIII_2_CIV_2_ on graphene oxide-coated grids (*left*) and holey gold grids (*right*). **D,** Three-dimensional reconstruction of mycobacterial CIII_2_CIV_2_ in its native membrane.

### Structure of mycobacterial CIII_2_CIV_2_ from native membranes

We next attempted a similar analysis for CIII_2_CIV_2_. While a number of CIII_2_CIV_2_ structures have been determined previously, all were obtained following detergent solubilization of the membrane (Gong et al., 2018; Wiseman et al., 2018; Yanofsky et al., 2021; S. Zhou et al., 2021). As with ATP synthase, we prepared IMVs, but this time from a strain with a 3×FLAG tag at the C terminus of the QcrB subunit of CIII_2_CIV_2_ encoded by the genomic DNA (Wiseman et al., 2018; Yanofsky et al., 2021). As done with ATP synthase, IMVs containing CIII_2_CIV_2_ were enriched by affinity chromatography and further purified by size exclusion chromatography. Owing to the higher yield of IMVs from this strain, cryo-EM specimens could be prepared either using graphene-oxide coated grids with diluted samples (**Fig. 1C**, *left*) or using holey gold coated grids with more concentrated samples (**Fig. 1C**, *right*). In both cases, the images appeared more heterogeneous than with the ATP synthase sample. However, there was less background noise in images from holey gold and these images were used for subsequent analysis. Unlike the ATP synthase that protrudes from the membrane, it was difficult to identify individual CIII_2_CIV_2_ complexes in the images. Instead, CIII_2_CIV_2_ complexes were located by exhaustive selection of lipid bilayer regions at the edges of vesicles in images. These regions were subjected to multiple rounds of 2D classification and the resulting set of particle images used to train a deep neural network particle selection algorithm (Bepler et al., 2019) (**Fig. S3**). Employing this strategy, it was possible to obtain 2D classes corresponding to the known structure of CIII_2_CIV_2_ (**Fig. 1C**, *lower*). Particle images from these classes were used to calculate a C2-symmetric map of the enzyme to a nominal overall resolution of 3.2 Å (**Fig. 1D**, **Fig. S4**, **Table S1**). Interestingly, while many particle images that contributed to the final 3D map came from intact IMVs (**Fig. S5A**, *green circles*), most came from flat patches of bilayer (**Fig. S5A**, *orange circles*). In the structure of CIII_2_CIV_2_, the patch of bilayer around the complex is mostly flat with a slight curvature toward the periplasmic side of the complex (**Fig. 1D**, **Fig. S5B**). This patch of nearly flat bilayer appears larger than can be accommodated by the ∼50 nm diameter IMVs, suggesting why many particles were found in patches of bilayer detached from the vesicles. The map fit an atomic model of CIII_2_CIV_2_ (Yanofsky et al., 2021) with high fidelity (**Fig. 2A**, **Fig. S5C**, **Table S2**), clearly resolving the two leaflets of the lipid bilayer (**Fig. 1D** and **2B**, *semi-transparent density*). Lipids and MQ molecules seen in maps of the detergent solubilized enzyme (Gong et al., 2018; Yanofsky et al., 2021) were also present in the map of the enzyme from the native lipid bilayer (**Fig. S5D**).

**Figure 2.**
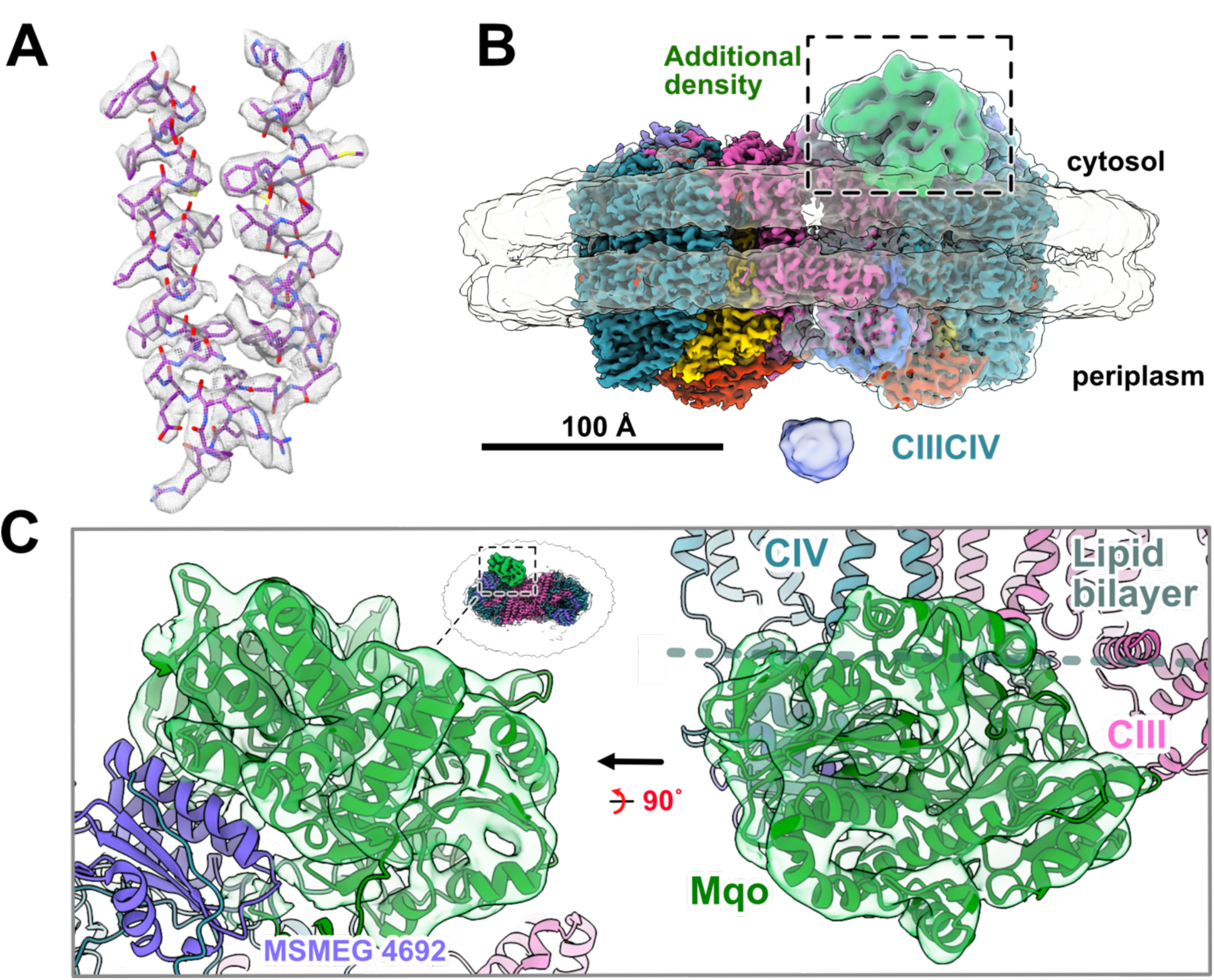
Interaction of Mqo with CIII_2_CIV_2_ in native membranes. **A,** Example high-resolution map region with fitted atomic model, corresponding to His110 to Phe157 of the QcrB subunit. **B,** Three-dimensional reconstruction showing an additional density attached to a CIIICIV complex from the CIII_2_CIV_2_ structure. A low-resolution reconstruction including the additional density is shown as a semi-transparent surface. **C,** Fitting of an atomic model for Mqo (*green*) into the additional map density.

### Malate:quinone oxidoreductase binds CIII_2_CIV_2_ in native membranes

In the C2-symmetric CIII_2_CIV_2_ map, faint additional densities were detected on the cytosolic side of the supercomplex. Classification focusing on this region of the map, followed by local refinement (**Fig. S3**), revealed an additional protein density at ∼6-9 Å resolution (**Fig. 2B**, **Fig. S4C** and **D**). This density was derived from ∼30% of the CIIICIV regions in the CIII_2_CIV_2_ particle images. The additional density had a volume corresponding to a protein of ∼50 kDa and included seven α-helix-like features and a small flat region consistent with a β-sheet. To identify the protein, we attempted to manually fit AlphaFold models into the density (Jumper et al., 2021), testing models of all *M. smegmatis* enzymes related to the ETC and central carbon metabolism (including enzymes from the Krebs cycle, glycolysis, and the pentose phosphate pathway). The AlphaFold model for Mqo fit the density unambiguously (**Fig. 2C** and **Fig. S5E**, *green density*). In the structure, Mqo is bound between the CIII and CIV portion of the CIII_2_CIV_2_ supercomplex and makes close contacts with the MSMEG_4692 subunit (**Fig. 2C**, *purple density*), a protein that did not have a known function in the enzyme. Part of Mqo appears to be embedded in the bilayer as suggested previously (Harold et al., 2022).

Mqo was not detected in maps of detergent-solubilized CIII_2_CIV_2_ (Gong et al., 2018; Wiseman et al., 2018; Yanofsky et al., 2021; S. Zhou et al., 2021). Similarly, Mqo was not identified in mass spectrometry of the purified CIII_2_CIV_2_ (Gong et al., 2018; Wiseman et al., 2018). Both these observations suggest that the Mqo:CIII_2_CIV_2_ interaction is easily and completely disrupted by detergent. To confirm detergent-induced dissociation of the interaction, we constructed an *M. smegmatis* strain with Mqo bearing a C-terminal 3×FLAG tag. Affinity purification from this strain following detergent solubilization of the membrane yielded pure Mqo, but did not show bands for CIII_2_CIV_2_ on an SDS-PAGE gel (**Fig. S5F**). In contrast, detergent-free affinity purification of IMVs from this *M. smegmatis* strain provided a high yield of vesicles (**Fig. S5G**), consistent with Mqo being attached to the IMV membrane. Together, these observations indicate that, as expected, Mqo behaves as a membrane protein and that the Mqo:CIII_2_CIV_2_ interface does not tolerate detergents. The finding that Mqo binds CIII_2_CIV_2_ in *M. smegmatis* provides an unexpected additional link between the Krebs cycle and the ETC.

### Fumarate hydratase class II provides malate to Mqo in IMVs

The binding of Mqo to CIII_2_CIV_2_ prompted us to investigate the previous unanticipated observation that fumarate, the precursor to malate in the Krebs cycle, can drive the ETC in mycobacterial IMVs (Courbon et al., 2023; Harden et al., 2024) (**Fig. 3A** and **Fig. 3B**, *black curve*). Fumarate is converted to malate by one of two *M. smegmatis* fumarate hydratases: fumarate hydratase class I (FH1, MSMEG_2985) and fumarate hydratase class II (FH2, MSMEG_5240). Experimental (Baugh et al., 2015) and predicted (Jumper et al., 2021) structures suggest that both FH1 and FH2 are soluble enzymes, making it surprising that this activity is found with purified IMVs. We hypothesized that one of the fumarate hydratases copurifies with IMVs, resulting in conversion of fumarate to malate at the membrane. Subsequent malate oxidation by Mqo would provide electrons to the ETC, which translocates protons into the IMV. To test this hypothesis, we first attempted to identify whether FH1 or FH2 is necessary for fumarate-driven IMV acidification. We prepared separate Δ*fh1* and Δ*fh2* deletion mutant strains and tested the ability of fumarate to drive acidification of IMVs prepared from the membrane fraction of these bacteria. IMVs from the Δ*fh1* strain showed robust fumarate-driven acidification (**Fig. 3B**, *orange curve*), comparable to IMVs from wildtype *M. smegmatis* (**Fig. 3B**, *black curve*). In contrast, IMVs from the Δ*fh2* strain showed negligible fumarate-driven acidification (**Fig. 3B**, *red curve*), indicating that the relevant fumarate hydratase activity was from FH2.

**Figure 3.**
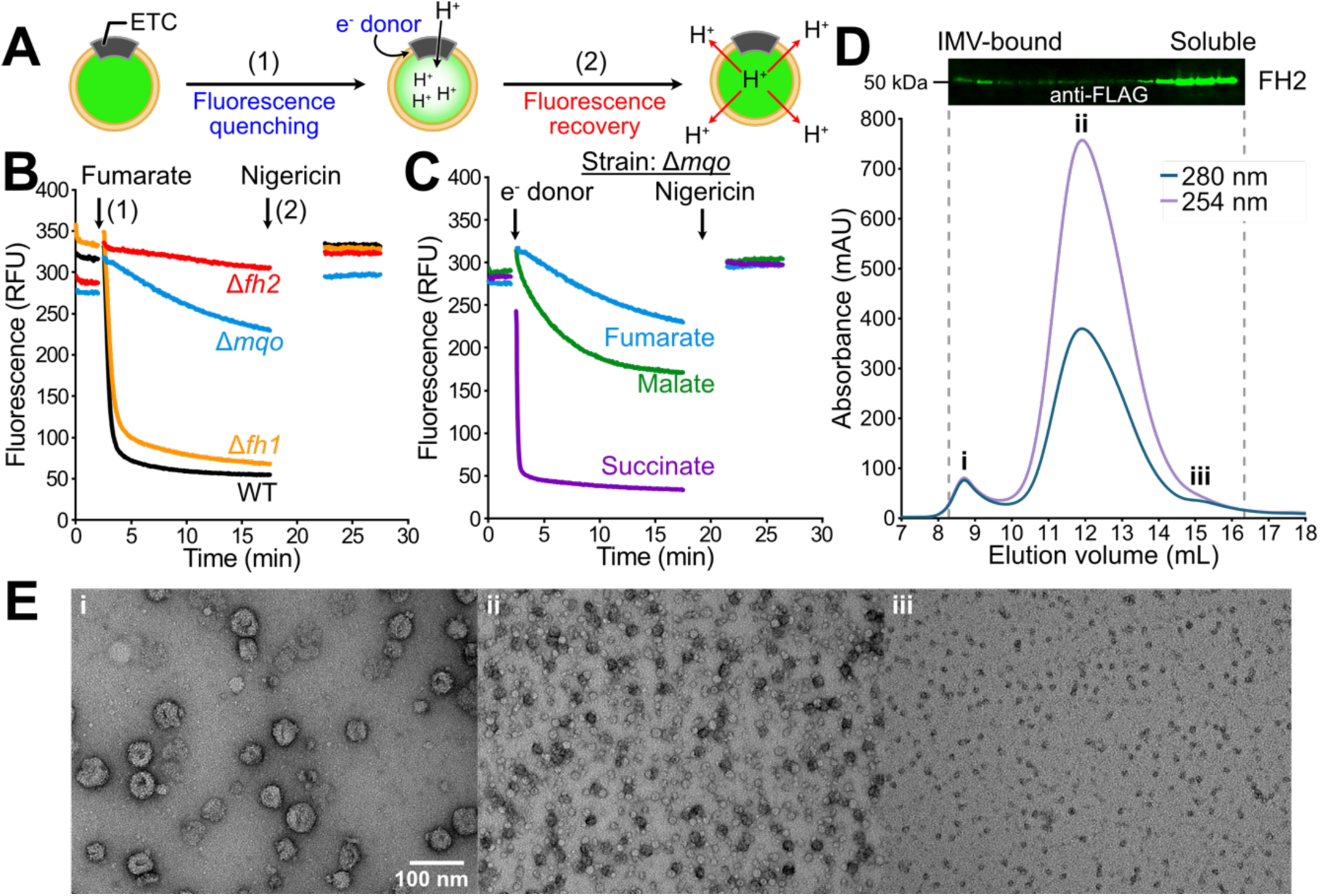
FH2 co-purifies with *M. smegmatis* IMVs. **A,** Schematic for the IMV acidification assay. Step (1): injection of an electron donor leads to IMV acidification and ACMA fluorescence quenching. Step (2): injection of nigericin collapses the proton motive force causing fluorescence recovery. **B,** Fumarate-induced IMV acidification with different knockout strains compared to a wildtype strain. RFU, relative fluorescence units. **C,** IMV acidification assay with the Δ*mqo* strain using various electron donors. **D,** Western blot against the FLAG epitope of FH2-3×FLAG in size exclusion chromatography fractions (*upper*). Size exclusion chromatogram of sample prepared from FH2-3×FLAG after affinity purification (*lower*). Roman numerals indicate different peaks. **E,** Negative stain electron micrographs from the different size exclusion peaks showing IMVs (i), ribosomes (ii), and FH2 (iii) purified with FH2-3×FLAG. Assay results are representative from two or more independent preparations of material.

We next tested the second part of the hypothesis, that FH2 provides Mqo with malate to drive IMV acidification. We prepared a Δ*mqo* deletion strain, isolated IMVs, and performed acidification assays. In support of the hypothesis, fumarate-driven acidification was greatly attenuated in these assays (**Fig. 3B**, *blue curve*) compared to wildtype IMVs (**Fig. 3B**, *black curve*). The residual activity is explained by the known ability of malate to serve as an electron donor for Complex II, which provides MQH_2_ to CIII_2_CIV_2_ (Vinogradov et al., 1989). To confirm this explanation, we tested the ability of IMVs from the Δ*mqo* strain to acidify with different electron donors. These experiments show that in addition to fumarate (**Fig. 3C**, *blue curve*), malate can also drive acidification of Δ*mqo* IMVs (**Fig. 3C**, *green curve*), although not as effectively as succinate (**Fig. 3C**, *purple curve*), which is the preferred substrate of Complex II. These observations support the model that oxidation of fumarate to malate at the IMV membrane provides substrate to Mqo.

To test whether FH2 associates with membranes, we prepared an *M. smegmatis* strain with a 3×FLAG at the C terminus of FH2 encoded by the genomic DNA. IMVs were isolated from this strain and subjected to anti-FLAG affinity chromatography and size exclusion chromatography (**Fig. 3D**, *lower*). The resulting chromatogram shows a small peak centered at 8.7 mL that negative stain electron microscopy (**Fig. 3E-i**) reveals were IMVs purified with the affinity-tagged FH2. A large peak at 12.0 mL in the chromatogram is the result of ribosomes that frequently contaminate mycobacterial IMV preparations, as seen from the strong absorbance at 254 nm and by negative stain electron microscopy (**Fig. 3E-ii**). Western blotting for the 3×FLAG on FH2 shows that the vast majority of FH2 elutes later than the IMVs and ribosomes (**Fig. 3D**, *upper*, **Fig. 3E-iii**). Together, these observations indicate that FH2 is a soluble enzyme but that it co-purifies with IMVs, probably through a weak non-specific interaction with the membrane. This interaction is consistent with mass spectrometry that identified FH2 in both the cytosolic and membrane fractions of *M. tuberculosis*, or entirely in the membrane fraction of cells (de Souza et al., 2011; Målen et al., 2010; Mattow et al., 2003; Mawuenyega et al., 2005; Xiong et al., 2005). The knowledge that FH2 co-purifies with IMVs is essential for interpreting experiments that probe ETC activity by providing succinate or fumarate as an electron donor.

### Disruption of CIII_2_CIV_2_ prevents malate-driven ETC activity

To investigate the association of Mqo with CIII_2_CIV_2_, we prepared IMVs from an *M. smegmatis* strain lacking the operon that encodes CIII from CIII_2_CIV_2_ (Δ*qcr*) (Chong et al., 2020). Using these Δ*qcr* IMVs we performed IMV acidification assays and measured oxygen consumption with different electron donors. When provided with NADH as an electron donor, the Δ*qcr* IMVs consumed oxygen at a rate comparable to IMVs from a wildtype strain (**Fig. 4A**, *right*), presumably through cyt. *bd*, which is upregulated upon disruption of CIII_2_CIV_2_ (Beites et al., 2019). In contrast, when malate was provided as the electron donor, the wildtype IMVs consumed oxygen but the Δ*qcr* IMVs did not (**Fig. 4A**, *left*). Similarly, IMVs from the Δ*qcr* strain displayed NADH-driven acidification (**Fig. 4B**, *orange*) but not malate-driven acidification (**Fig. 4B**, *green*). These observations indicate that Mqo requires CIII_2_CIV_2_ to feed electrons into the mycobacterial ETC.

**Figure 4.**
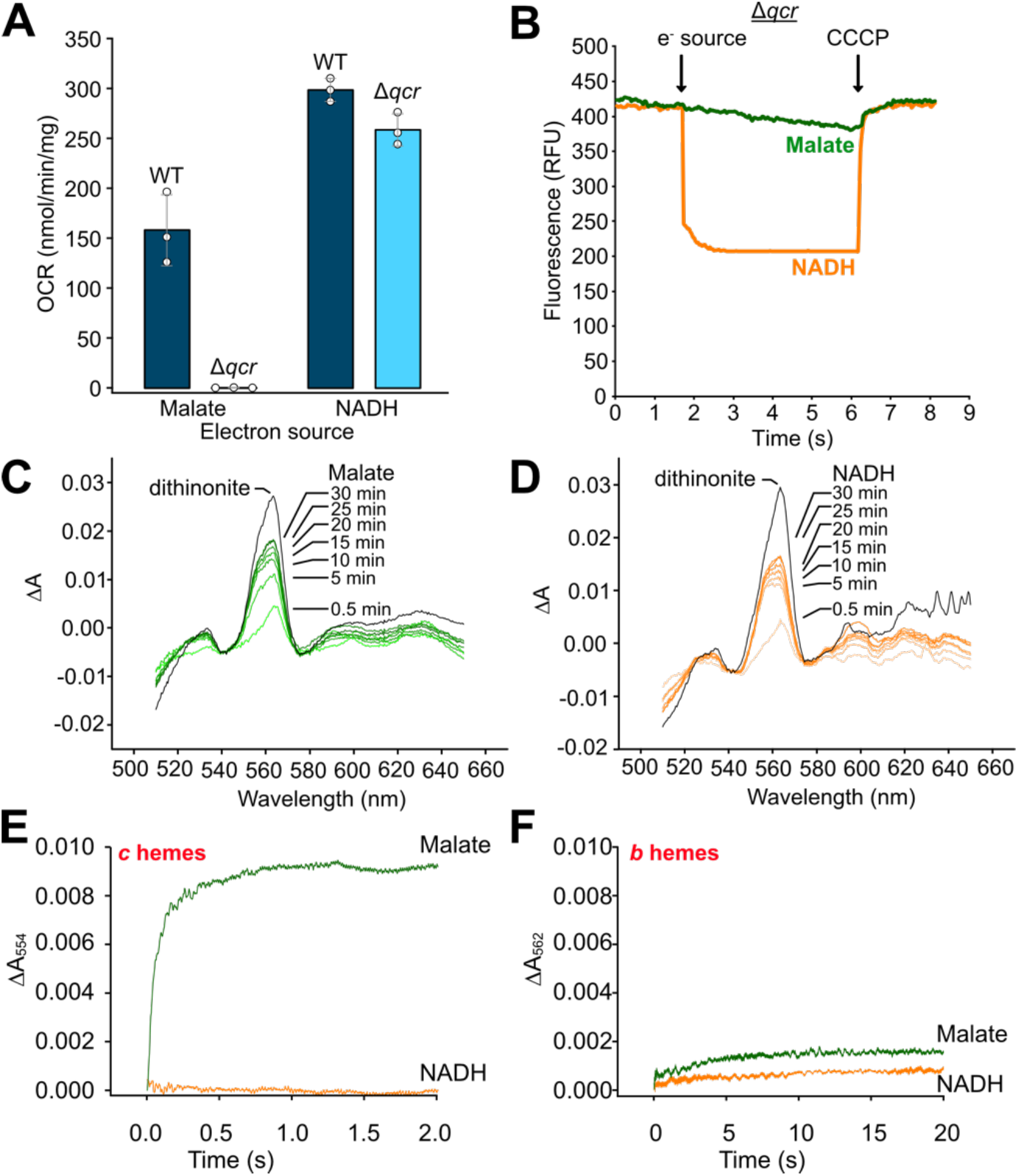
Rapid electron transfer from malate to CIII_2_CIV_2_. **A,** Oxygen consumption rate (OCR) for IMVs from wildtype (WT) and Δ*qcr M. smegmatis* when provided with malate or NADH. Mean and three technical replicates from a representative biological replicate are shown. **B,** IMV acidification assay for IMVs from wildtype (WT) and Δ*qcr M. smegmatis* when provided with malate or NADH. IMV assays are representative of three technical replicates. **C,** Reduced minus oxidized difference spectra at different time points for IMVs reduced with malate. Complete reduction was achieved by addition of dithionite. **D,** Reduced minus oxidized difference spectra over time for IMVs reduced with NADH. Complete reduction was achieved by addition of dithionite. **E,** Stopped-flow spectroscopy measuring the change in absorbance at 554 nm (ΔA_554_), corresponding to reduction of *b* hemes, on addition of malate and NADH. **F,** Stopped-flow spectroscopy measuring the change in absorbance at 562 nm (ΔA_562_), corresponding to reduction of *c* hemes, on addition of malate and NADH. Spectroscopic experiments are representative from two independent preparations of IMVs.

### Association of Mqo with CIII_2_CIV_2_ enables rapid electron transfer from Mqo to CIII_2_CIV_2_

The physical association of Mqo with CIII_2_CIV_2_ suggests the possibility of electron channeling from one to the other. We performed optical spectroscopy of IMVs to investigate reduction of the different hemes in the sample. In these experiments, oxidation of CIV (cyt. *aa*_3_) was inhibited by addition of potassium cyanide and the sample was subsequently reduced by addition of malate or NADH. Malate transfers electrons to the ETC via Mqo while NADH transfers electrons via NDH-2. Reduction of the sample with succinate was avoided because of the confounding effect that succinate is converted to fumarate by Complex II, which is in turn converted to malate by FH2. We performed initial experiments with a conventional spectrophotometer. Spectra of an oxidized sample were subtracted from spectra of reduced samples, with increased absorbance at 554 nm (ΔA_554_) corresponding to reduction of *c* hemes, and increased absorbance at 562 nm (ΔA_562_) corresponding to reduction of *b* hemes. Addition of malate (**Fig. 4C**) led to a ΔA_554_ of 0.0098 over ∼30 min, which increased to 0.013 on complete reduction with dithionite, indicating that malate can reduce ∼75% of the *c* hemes in the sample. Addition of malate also led to a ΔA_562_ of 0.018 over ∼30 min, which increased to 0.027 with addition of dithionite, indicating malate reduces ∼67% of the *b* hemes in the sample. Similarly, addition of NADH (**Fig. 4D**) led to a ΔA_554_ of 0.0076 and ΔA_562_ of 0.0165 over ∼30 min, which increased to 0.011 and 0.0300, respectively, on complete reduction with dithionite, indicating that NADH reduces ∼69% of the *c* hemes and ∼55% of the *b* hemes in the sample. Negligible change in absorbance at 630 nm (*d* hemes) or 603 nm (*a* hemes) was observed on addition of malate or NADH. Lack of reduction of *d* hemes suggests that the *b* hemes detected are from CIII_2_CIV_2_ (cyt. *bcc*-*aa_3_*) not cyt. *bd*. This absence of signal from cyt. *bd* was expected because of the low level of this complex in the sample, as evident from the observation that the CIII_2_CIV_2_-specific inhibitor Q203 can completely block succinate- and NADH-driven acidification of wildtype IMVs (Harden et al., 2024). Reduction of heme *a*_3_ was likely not observed because of cyanide binding, while the midpoint potential of heme *a* is probably not high enough (compared to that of the *c* hemes) to allow its reduction when further electron transfer to heme *a*_3_ and the associated Cu_B_ is blocked.

In contrast with experiments over long timescales in a conventional spectrophotometer, stopped-flow spectroscopy showed that addition of malate caused reduction of the *c* hemes (ΔA_554_) within ∼500 msec (**Fig. 4E**, *green curve*) while addition of NADH (**Fig. 4E**, *orange curve*) did not reduce *c* hemes on this timescale. The ΔA_554_ of ∼0.009 indicates that ∼30% of the *c* hemes are rapidly reduced on addition of malate, which matches the ∼30% of CIII complexes that were observed to have a bound Mqo by cryo-EM (88,444 CIIICIV regions out of 293,585 from 146,585 particle images). Measurement of ΔA_562_ indicates that the *b* hemes are not reduced rapidly by either malate or NADH (**Fig. 4F**, *orange and green curves*). Therefore, the comparison of conventional spectroscopy and stopped-flow spectroscopy shows that electrons are transferred from malate to CIII_2_CIV_2_ much faster than from NADH to CIII_2_CIV_2_, consistent with rapid transfer of electrons within the Mqo:CIII_2_CIV_2_ complex.

## Discussion

In this study we show that it is possible to obtain high-resolution structures of membrane proteins in their native membranes even when they are not the major species in the lipid bilayer. While the complexes analyzed here are larger than many membrane proteins, further advances in imaging and image analysis could allow for structure determination of other membrane proteins in their native membrane. These structures would likely allow for identification of other protein-protein interactions that have been missed previously.

Imaging of mycobacterial IMVs shows that Mqo is a subunit of the CIII_2_CIV_2_ supercomplex, which allows rapid channeling of electrons from malate to the *c* hemes of the assembly. Complex III within CIII_2_CIV_2_ contributes to the pmf through a mechanism known as a “Q cycle” (Sarewicz and Osyczka, 2015) (**Fig. 5A**). Briefly, oxidation of MQH_2_ at the Q_P_ site releases two protons to the positive (periplasmic) side of the membrane. The first electron is transferred sequentially to the iron-sulfur (FeS) center, heme *c*_I_ and *c*_II_ from the cyt. *cc* domain, and the binuclear Cu_A_ of Complex IV. The second electron is transferred to hemes *b_L_* and then *b*_H_, before reducing MQ in the Q_N_ site of the enzyme. Oxidation of two MQH_2_ molecules in the Q_P_ site results in full reduction of MQ in the Q_N_ site and abstraction of two protons from the negative (cytosolic) side of the membrane. The experiments described here show that malate oxidation by

**Figure 5.**
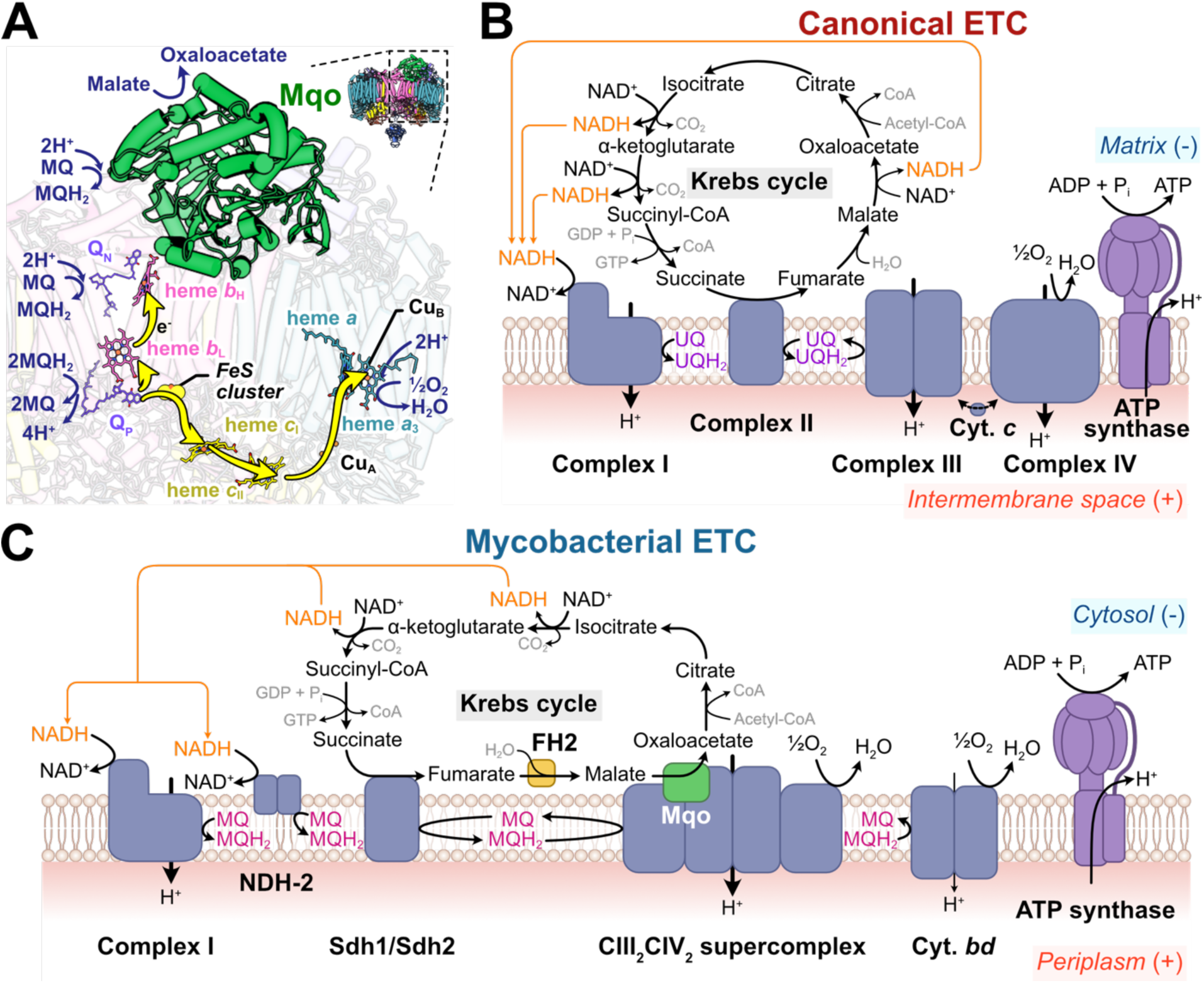
Mechanism and role of Mqo in the mycobacterial ETC. **A,** The Q cycle in CIII with Mqo bound nearby. **B,** Canonical Krebs cycle and ETC. **C,** Mycobacterial Krebs cycle and ETC.

Mqo rapidly reduces the *c* hemes but not the *b* hemes in CIII, even though the *b* hemes are closer to Mqo than the *c* hemes (**Fig. 5A**), and the *b* hemes are reduced before the *c* hemes in the Q cycle. There are two possible explanations for this observation. First, previous structures of CIII_2_CIV_2_ have shown MQ bound in the Q_N_ site (Gong et al., 2018; Wiseman et al., 2018; Yanofsky et al., 2021) and an excess of oxidized MQ is expected in the membrane. This MQ would rapidly oxidize the *b* hemes and prevent observation of *b* heme reduction in the stopped flow experiments described here. In contrast, rapid oxidation of the *c* hemes is prevented by blocking downstream electron transfer to CIV with cyanide. Further, the *b* hemes have the lowest midpoint potential of all of the hemes in the obligate CIII_2_CIV_2_ supercomplex of actinobacteria (Kao et al., 2016). Therefore, in the unlikely event that all the MQ in the membrane was reduced faster than the time resolution of the stopped flow experiment, even if electrons were transferred via the *b* hemes, one would not see reduction of the *b* hemes until the *c* hemes were reduced. It remains unclear whether electrons from Mqo are transferred first to a bound MQ or directly to the hemes of CIII.

The canonical Krebs cycle and ETC are linked by the transfer of NADH from the Krebs cycle to Complex I of the ETC and the shared activity of Complex II in the two processes (**Fig. 5B**). In contrast, both Complex II and Mqo:CIII_2_CIV_2_ are shared between the Krebs cycle and ETC in mycobacteria, with two acceptors of NADH from the Krebs cycle in the ETC (**Fig. 5C**). Both Mqo and CIII_2_CIV_2_ are essential for *M. smegmatis* growth on non-fermentable carbon sources in minimal medium (Harold et al., 2022), consistent with a physical organization that ensures that carbon sources metabolized by the Krebs cycle drive respiration via CIII_2_CIV_2_. Both Mqo and MDH appear to be important for survival of *M. tuberculosis* (Kumar et al., 2024) as well as other actinobacteria where both enzymes are present (Molenaar et al., 1998). The link between Mqo and CIII_2_CIV_2_ explains why Mqo and MDH are not equivalent. The physical association of the enzymes of the Krebs cycle and ETC is reminiscent of the idea of a metabolon, a physical grouping of enzymes of the Krebs cycle for substrate shuttling. This idea, which has persisted for decades (Srere, 1985), remains controversial (Omini et al., 2024). However, mycobacteria have been demonstrated to possess compartmentalized catabolism, using different metabolic pathways to selectively consume different carbon sources (de Carvalho et al., 2010). The experiments shown here suggest the possibility of bioenergetic compartmentalization, with different pathways feeding into different branches of the mycobacterial ETC.

## Methods

### Preparation of M. smegmatis strains

*M. smegmatis* strains were generated using the ORBIT method (Murphy et al., 2018). In this approach, the parent strain is transformed with a plasmid encoding the Che9c phage RecT annealase and Bxb1 integrase, a payload plasmid with the desired insert, and an oligonucleotide that guides integration of the payload into the chromosomal DNA. The resulting strains were selected on LB plates with hygromycin (50 μg/mL) and for deletion strains plates were supplemented with 20 mM glucose. Addition of the 3×FLAG sequence used the plasmid pSAB41 (Guo et al., 2021) as the payload plasmid while gene deletion used the plasmid pKM464 (Addgene plasmid #108322) as the payload plasmid. Insertion of the 3×FLAG sequence, or deletion of a gene, was confirmed by colony PCR. The targeting oligonucleotide sequences were gtaaagcgttggcgcatcgaggtctttggcccagctgcggcgcagtgacaccgtcatgtgtctcccgccgtcaGGTTTGTACCGTACACCACTGAGACCGCGGTGGTTGACCAGACAAACCcacggtcgccggggcgttggctgcttccgctgt attcgcttgcacgtccagcttgagcaccttggtgccc (addition of 3×FLAG tag to Mqo, MSMEG_2613, Strain SABM15), ccagctgcggcgcagtgacaccgtcatgtgtctcccgccgtcacacggtcgccggggcgttggctgcttcGGTTTGTACCGTACACCACTGAGACCGCGGTGGTTGACCAGACAAACCcgcgttcgtacctgcgtttgcctcagacaccctgatcgctagcccttcttcgtcgcgtggcccatgaacc (deletion of Mqo, MSMEG_2613, strain SABM18), agggcatgctgttcgatctggccgcacggatcgaacaccgatgaccgtcgactacacaccgatactgccgGGTTTGTCTGGTCAACCACCGCGGTCTCAGTGGTGTACGGTACAAACCccactactcacgatcggagcagcaccgcagtgattctcacgcacgcagagccacttcacctcagcctgga (deletion of FH1, MSMEG_2985, strain SABM20), caagctgagcctcgaggaactcgaccgtcgtctcgacgtgctcgccatggcgcgggtcaaggacggcgagGGTTTGTCTGGTCAACCACCGCGGTCTCAGTGGTGTACGGTACAAACCtaggggtcaccttcctctcgcgacttcagtcggggagcggcgcggtccagcgcaggatcgtgccgccgcccgg (addition of 3×FLAG tag to FH2, MSMEG_5240, strain SABM16), ctgccgtaacacaccaacaaacaagggacgcaacagcacaatggccgacaccgacgtcgaataccgcatcGGTTTGTCTGGTCAACCACCGCGGTCTCAGTGGTGTACGGTACAAACCctcgccatggcgcgggtcaaggacggcgagtaggggtcaccttcctctcgcgacttcagtcggggagcggcgc (deletion of FH2, MSMEG_5240, strain SABM17).

### Preparation of ATP synthase-enriched IMVs

The previously engineered *M. smegmatis* strain GMC_MSM1, with a C-terminal 3×FLAG tag on the β subunit of ATP synthase (Guo et al., 2021), was grown for two days at 30 °C on LB plates supplemented with 50 µg/mL hygromycin. Bacteria from the plate were transferred to a 25 mL preculture of 7H9 ADS medium (7H9 from Sigma with 5 g/L albumin, 2 g/L dextrose, 0.8 g/L NaCl) and grown for 48 h in the dark at 30 °C with shaking at 180 rpm. This preculture was used to inoculate a 6 L culture in 7H9 TDS (7H9 with 10 g/L tryptone, 2 g/L dextrose, 0.8 g/L NaCl) and grown for 48 h in the dark. Bacteria were harvested by centrifugation at 6500 ×g for 20 min at 4 °C. Cell pellets were resuspended in up to 150 mL lysis buffer (50 mM Tris-HCl pH 7.5, 150 mM NaCl, 5 mM MgSO_4_, 5 mM 6-aminocaproic acid, 5 mM benzamidine hydrochloride) and frozen for further use. Thawed cells were passed through an Avestin homogenizer three times at 20 kpsi and debris was removed by centrifugation at 39000 ×g for 30 min. Membrane pellets were collected by centrifugation at 200,000 ×g (Beckmann 70 Ti rotor) and resuspended in 40 mL IMV S-buffer (50 mM Tris-HCl pH 7.5, 150 mM NaCl, 20% glycerol, 5 mM MgSO_4_, 5 mM benzamidine hydrochloride, 5 mM 6-aminocaproic acid, 1 mM PMSF, 25 µg/ml DNaseI). IMVs were filtered with a 0.45 μm filter (Corning) and loaded on a 2 mL column of M2 affinity matrix (Sigma) equilibrated with IMV S-buffer. After washing with IMV S-buffer, bound proteins were eluted with three column volumes of IMV S-buffer with 150 μg/mL 3×FLAG peptide. Purified IMVs were concentrated with a 100 kDa concentrator and loaded on a Superose 6 Increase 10/300 GL gel filtration column (Cytiva) equilibrated with IMV SEC buffer (50 mM Tris-HCl pH 7.5, 150 mM NaCl, 15% glycerol, 5 mM MgCl_2_, 5 mM benzamidine hydrochloride, 5 mM 6-aminocaproic acid). IMVs were further concentrated, and buffer exchanged with a Zeba spin desalting column (Thermo Scientific) equilibrated with TBS buffer (50 mM Tris-HCl pH 7.5, 150 mM NaCl).

### Preparation of CIII_2_CIV_2_-enriched IMVs

A previously engineered *M. smegmatis* strain with a C-terminal 3×FLAG tag on QcrB (Yanofsky et al., 2021) was grown on 7H9 plates with hygromycin for 48 h at 30 °C. A colony from the plate was transferred to a 20 mL LB preculture with 50 µg/mL hygromycin and grown for 48 h in the dark at 30 °C and 180 rpm shaking, which was used to inoculate a 6 L culture in 7H9 TDS and grown for 48 h in the dark. Cells were harvested by centrifugation at 6500 ×g for 20 min at 4 °C, and then cell pellets were frozen in liquid nitrogen and stored at −80 °C. Thawed cell pellets were resuspended in ∼150 mL lysis buffer (50 mM Tris-HCl pH 7.5, 100 mM NaCl, 0.5 mM EDTA) and homogenized first with a Dounce homogenizer, then filtered with cheesecloth and passed through an Avestin homogenizer three times at 20 kpsi. Lysed cells were centrifuged at 39000 ×g for 30 min to remove cell debris. The supernatant was then centrifuged at 149,000 ×g for 60 min (Beckmann 70 Ti rotor) to collect the membranes. Membrane pellets were homogenized in IMV buffer (50 mM Tris-HCl pH 7.5, 150 mM NaCl, 5 mM MgSO_4_; 12 mL/g membranes) with a Dounce homogenizer before aliquoting in falcon tubes, freezing in liquid nitrogen, and storing at −80 °C. IMVs were thawed and loaded onto 0.5-1.5 mL of M2 resin, which was equilibrated with IMV buffer. The column was then washed with ∼5 column volumes of IMV buffer before elution with ∼8 column volumes of 150 μg/mL 3×FLAG peptide in IMV buffer. The eluant was concentrated to ∼150 μL and loaded onto a Superose 6 Increase 10/300 GL gel filtration column previously equilibrated with IMV buffer. Fractions containing signal at 420 nm, which corresponds, in part, to absorbance from hemes were collected.

### Preparation of FH2-enriched IMVs, Mqo-enriched IMVs, and detergent solubilized Mqo

Purification of IMVs from the *M. smegmatis* strain containing a C-terminal 3×FLAG on FH2 followed the same procedure as the purification of ATP synthase-enriched IMVs, with the following exceptions: the preculture was started in 25 mL 7H9 TDS, 1 mL of M2 affinity matrix was used, and the gel filtration column was equilibrated using a SEC buffer containing 50 mM Tris-HCl pH 7.4 (at 4 °C), 150 mM NaCl, 20% glycerol, and 5 mM MgSO_4_. For Mqo-enriched IMVs the protocol was the same as for CIII_2_CIV_2_-enriched IMVs. To purify Mqo, membranes were solubilized in 1% dodecyl maltoside (DDM) for 60 min before removing the insoluble fraction by ultracentrifugation at 149,000 ×g for 60 min. DDM (0.05 % (w/v)) was included in all subsequent buffers.

### Freezing of cryo-EM grids

For freezing CIII_2_CIV_2_ IMVs on holey gold grids, the sample was concentrated until the absorbance at 420 nm was ∼0.4, which was empirically determined to be the optimal concentration. IMV sample (2 μL) was applied to nanofabricated holey gold grids (Marr et al., 2014) that were glow discharged for 2 min in air (20 mA with a PELCO easiGlow, Ted Pella) immediately prior to use. Grids were blotted for 2 s with an EM GP2 Plunge Freezing device (Leica Microsystems) held at 4 °C and 90% humidity before freezing in liquid ethane. When freezing ATP synthase IMVs or CIII_2_CIV_2_ IMVs on graphene oxide grids, the grids were glow discharged in air for 5 s with the grid bar side facing up before applying 5 µL of sample on the grid bar side of the grid. The grids were incubated for 3 min in the humidity chamber of the Leica EM GP2 plunge freezer at 4°C and 90% humidity, before removal of 3 µL of the sample with a pipettor, blotting for 5 s, and plunge freezing in liquid ethane.

### Cryo-EM data collection

Grids were screened with a Glacios 2 electron microscope (Thermo Fisher Scientific) operated at 200 kV and equipped with a Falcon 4i camera. A total exposure of ∼40 e^-^/Å^2^, exposure rate of ∼11.3 e^-^/pixel/s, and a nominal magnification of 92,000× with a calibrated pixel size of 1.5 Å were used. High-resolution data were collected with a Titan Krios G3 electron microscope (Thermo Fisher Scientific) operated at 300 kV and equipped with a Falcon 4i camera. Data collection was automated with the EPU software package.

### Image analysis for ATP synthase-enriched IMVs

All image analysis was performed in cryoSPARC. Particles were first manually picked to train an initial Topaz model. Particle images were then extracted using a 384 pixel box and curated using 2D classification. An initial 3D volume was then obtained by ab-initio reconstruction followed by homogenous or non-uniform refinement. This volume was used to subtract the membrane with a low-resolution mask around the ATP synthase. Subtracted particle images were then subjected to multiple rounds of ab-initio reconstruction and heterogenous refinement cleaning to obtain a rough ATP synthase structure. This approach was repeated several times using topaz models trained on the best particles from either 2D classification or 3D heterogenous refinement from the previous iteration. The best particle images from each iteration were then pooled, duplicates removed, and curated using the same approach as described above to obtain an initial structure of the ATP synthase (6,080 particle images). 3D classification and heterogenous refinement were then used to select particle images with a well resolved peripheral stalk (3,192 particle images). Subtracted particle images were re-extracted, and local refinement was performed to obtain a 9 Å resolution of the full ATP synthase in state 1, as well as an 8.2 Å resolution of the F_1_ region.

### Image analysis for CIII_2_CIV_2_-enriched IMVs

All image analysis was performed in cryoSPARC (Punjani et al., 2017). Particle images were extracted in 364-pixel boxes and all refinements performed were non-uniform refinements unless stated otherwise (Punjani et al., 2020). First, regions of bilayer were manually picked and used for template picking and picked particles were extracted. The extracted particle images were subject to 2D classification, and particle images in a class showing CIII_2_CIV_2_ (494 particle images) were used to train a Topaz model (Bepler et al., 2019). Duplicate particles were removed using a 120 Å minimum separation distance and 439,882 particle images were extracted and used to calculate 2D classes. Particles contributing to classes corresponding to CIII_2_CIV_2_ (29,204 particle images) were used in an ab-initio reconstruction with two classes. The particle images from the class corresponding to CIII_2_CIV_2_ (15,802 particle images) were used to train another Topaz model, with the particle images from this model added to the previous particle images. Duplicates were removed, and particle images were re-extracted. These particle images were once again used in an ab-initio reconstruction with several initial classes to clean out ‘junk particles’. The remaining particle images were again used to train a Topaz model, and the processes of extracting, removing duplicates, cleaning out junk, and training a Topaz model were repeated until refinement resolution and number of particle images remained unchanged between iterations. Reference-based motion correction, and per particle defocus correction and refinement of the 146,585 particle images with C2 symmetry led to a map of CIII_2_CIV_2_ at 3.2 Å resolution. Particle images were symmetry expanded and 3D variability analysis (Punjani and Fleet, 2021) and cluster analysis were performed with a low-pass filter of 8 Å and a generous mask placed over the cytosolic side of CIII_2_CIV_2_ where extra density was observed. The subpopulation of 88,444 particle images with extra density in the masked Mqo region were subject to local refinement with a mask over half of the supercomplex and Mqo. This refinement reached a nominal resolution of 3.4 Å and the final map was locally filtered to improve the density of the Mqo region, which was at a lower resolution then the rest of the map.

### Western blotting for FH2

From the 300 µL fractions following size exclusion chromatography of affinity-purified FH2-3×FLAG, alternating fractions from 8.2 to 16.3 mL were used for western blot analysis. Samples were subjected to SDS-PAGE, followed by semi-dry transfer onto a 0.45 µm nitrocellulose membrane. The primary antibody was anti-FLAG (F3165, Sigma) and the secondary antibody was fluorescent goat anti-mouse (LICOR), and both were used according to the manufacturer’s instructions. The membrane was imaged with an Odyssey^®^ XF Imaging System (LICOR) using the 800 nm channel for 30 seconds.

### Negative stain electron microscopy of IMVs

Continuous carbon film coated EM grids were glow discharged at 25 mA for 15 s with a PELCO easiGlow glow discharge device. IMV sample (3 µL) was applied, incubated for 2 min, and washed with three drops of water before staining with 2% (w/v) uranyl acetate for 1 min. The grid was then blotted from the edge and allowed to dry. Imaging was performed with a Hitachi HT7800 transmission electron microscope operating at 120 kV and equipped with the Xarosa 20 Megapixel CMOS Camera (EMSIS). Images were obtained at 50,000× magnification, corresponding to 2.93 Å/pixel, and with 4 µm defocus.

### Measurement of heme reduction

Electron transfer kinetics between malate or NADH and CIII_2_CIV_2_ were measured with a stopped-flow apparatus (Applied Photophysics). Fully oxidized IMVs (previously filtered with a Corning 0.45 μm syringe filter) in 50 mM Tris-HCl pH 7.5, 150 mM NaCl, 5 mM MgSO_4_ and 10 mM KCN were bubbled with N_2_ to minimize the amount of oxygen. The sample was then mixed with 500 µM malic acid or NADH in a 1:1 ratio. Reduction of CIII_2_CIV_2_ was monitored by the increase of absorbance at 554 nm (heme *c*), 562 nm (heme *b*), 603 nm (hemes *a* and *a*_3_), and 630 nm (heme *d*). For measurement of heme reduction over long timescales, optical absorption spectra were recorded with a Cary 100 UV-Vis spectrophotometer (Agilent Technologies) using the same buffer as for the stopped-flow experiments. Reduction of the hemes was observed by adding malic acid or NADH to a final concentration of 500 µM with a Hamilton syringe and measurements were taken every 5 min for 30 min. For the final measurement, a small aliquot of sodium dithionite was added to reduce the sample completely. The percentage of reduction for each time step was calculated assuming that the reduction obtained with sodium dithionite was complete.

### IMV acidification and oxygen consumption assays

IMV acidification experiments were performed in 96-well plates (BRANDplates™ pureGrade 96-well black microplates). Each reaction well consisted of 80 μL 2× ACMA assay buffer (20 mM HEPES-KOH pH 7.5, 200 mM KCl, 10 mM MgCl_2_), 10 μL of IMVs, 0.25 μL of 2 mM 9-amino-6-chloro-2-methoxyacridine (ACMA, dissolved in ethanol), and 26.55 μL of MilliQ water. Separate master mixes containing 1.5× the total volume needed were made in Eppendorf tubes for each condition and were kept on ice in the dark. After thorough mixing, 116.8 µL of the master mix solutions was transferred to each well of the 96-well plate. Fluorescence was measured at λ_ex_ = 410 nm and λ_em_ = 480 nm, at 25 °C, using a Synergy Neo2 HTS Hybrid Multi-Mode Plate Reader (BioTek). The 96-well plate was pre-incubated inside the plate reader for 5 min before fluorescence was monitored and recorded at 4 s intervals. Baseline readings of the samples were measured for 2 min, followed by automated injection of 40 µL electron donor (stock solutions of either 20 mM fumaric acid in 40 mM Tris, 20 mM malic acid in 40 mM Tris, or 20 mM sodium succinate in water) to each well to initiate proton pumping. The resulting final electron donor concentration was 5 mM in each well. Fluorescence was then measured for 15 min before 3.2 µL of 100 μM nigericin in 1% (v/v) ethanol was manually injected into each well with a multichannel pipettor and mixed with a different multichannel pipettor of larger volume. Fluorescence was then measured for an additional 5 min to observe recovery.

Analysis of fluorescence quenching for IMVs where the *qcr* operon was deleted was performed in a similar way with the following exceptions. IMVs were prepared from *M. smegmatis* cells grown under shaking (200 rpm) in Hartmans de Bont HdB minimal medium supplemented with 0.2% glucose, but without the specific addition of surfactants (i.e. tween 80 or tyloxapol). The fluorophore used was acridine orange (10 µM), proton pumping was initiated by the addition of either NADH (500 µM) or malic acid (5 mM), and reversal was achieved with 1 µM CCCP. Experiments are representative of a technical triplicate. Oxygen consumption rate (OCR) for IMVs was measured with an Oroboros Oxygraph-2k. Oxygen consumption was initiated with either 5 mM malate (final concentration) or 0.5 mM NADH as the sole electron donor.

## Author Contributions

JDT prepared specimens of the CIII_2_CIV_2_-enriched IMVs, and performed cryo-EM and image analysis for this sample. JY characterized fumarate and malate driven acidification of vesicles from different strains, the co-purification of FH2 with membranes, and prepared IMVs for spectroscopic analysis. GM Courbon prepared ATP synthase-enriched IMVs, imaged them, and performed image analysis. APLR performed spectroscopy and stopped-flow spectroscopy with IMVs. C-YC characterized acidification and oxygen consumption by Δ*qcr* IMVs. YL and CEC helped develop the methods for cryo-EM of membrane vesicles. SAB prepared Δ*fh1*, Δ*fh2*, and Δ*mqo M. smegmatis* strains. GM Cook planned and supervised experiments with Δ*qcr* IMVs. PB planned and supervised spectroscopic experiments and guided interpretation of results. JLR conceived the study, supervised and coordinated experiments, and coordinated the writing of the manuscript and preparation of the figures with the other authors.

## Data Deposition

The electron cryomicroscopy map and associated models described in this article have been deposited in the Electron Microscopy Data Bank (EMDB) with accession numbers EMD-47123 and EMD-47124 (ATP synthase maps) and EMD-46995 (Mqo:CIII_2_CIV_2_ map) and the Protein Data Bank with accession numbers PDB ID: 9DM1(Mqo:CIII_2_CIV_2_ model).

## Acknowledgements

We thank Masahiro Enomoto for teaching us how to make graphene oxide coated electron microscopy grids and Agnes Moe for supervision of the spectroscopic measurements at an early stage of the project. This work was supported by the Canadian Institutes of Health Research grant PJT162186 (JLR), grants from the Knut and Alice Wallenberg Foundation (2019.0043) and the Swedish Research Council (2022-03356) (PB), and Marsden Fund, Royal Society of New Zealand (GM Cook). JDT and CEC were supported by Canadian Institutes of Health Research (CIHR) Postdoctoral Fellowships, GM Courbon was supported by a Mary H. Beatty Fellowship, YL was supported by a CIHR Doctoral Canada Graduate Scholarship, and JLR was supported by the Canada Research Chairs program. Cryo-EM data were collected at the Toronto High-Resolution High-Throughput CryoEM Facility supported by the Canada Foundation for Innovation and Ontario Research Fund, ACMA assays were performed at the Structural and Biophysical Core at the Hospital for Sick Children.

## Supplementary Figure

**Figure S1.**
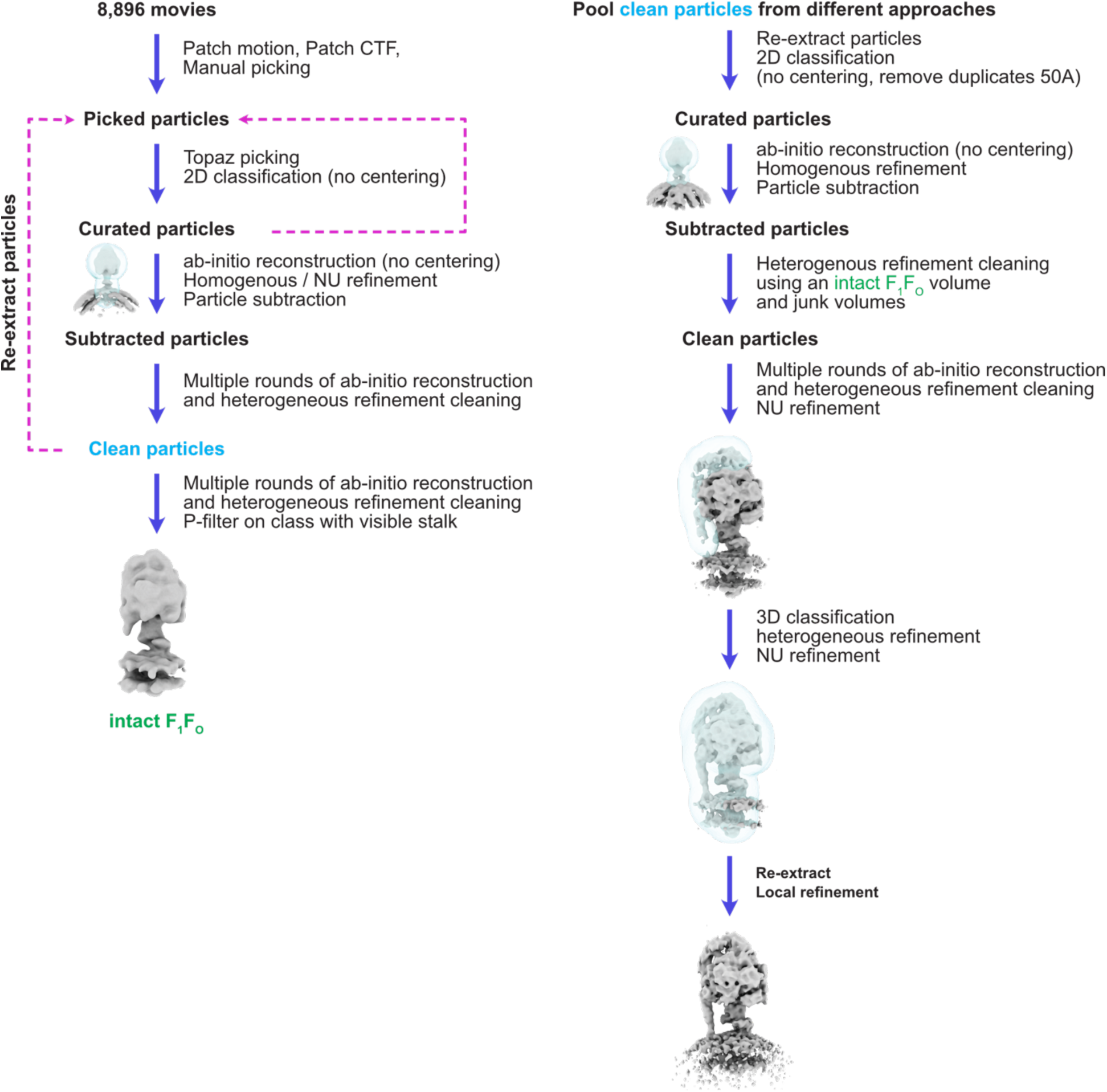
Workflow for calculating map of ATP synthase from IMV images.

**Figure S2.**
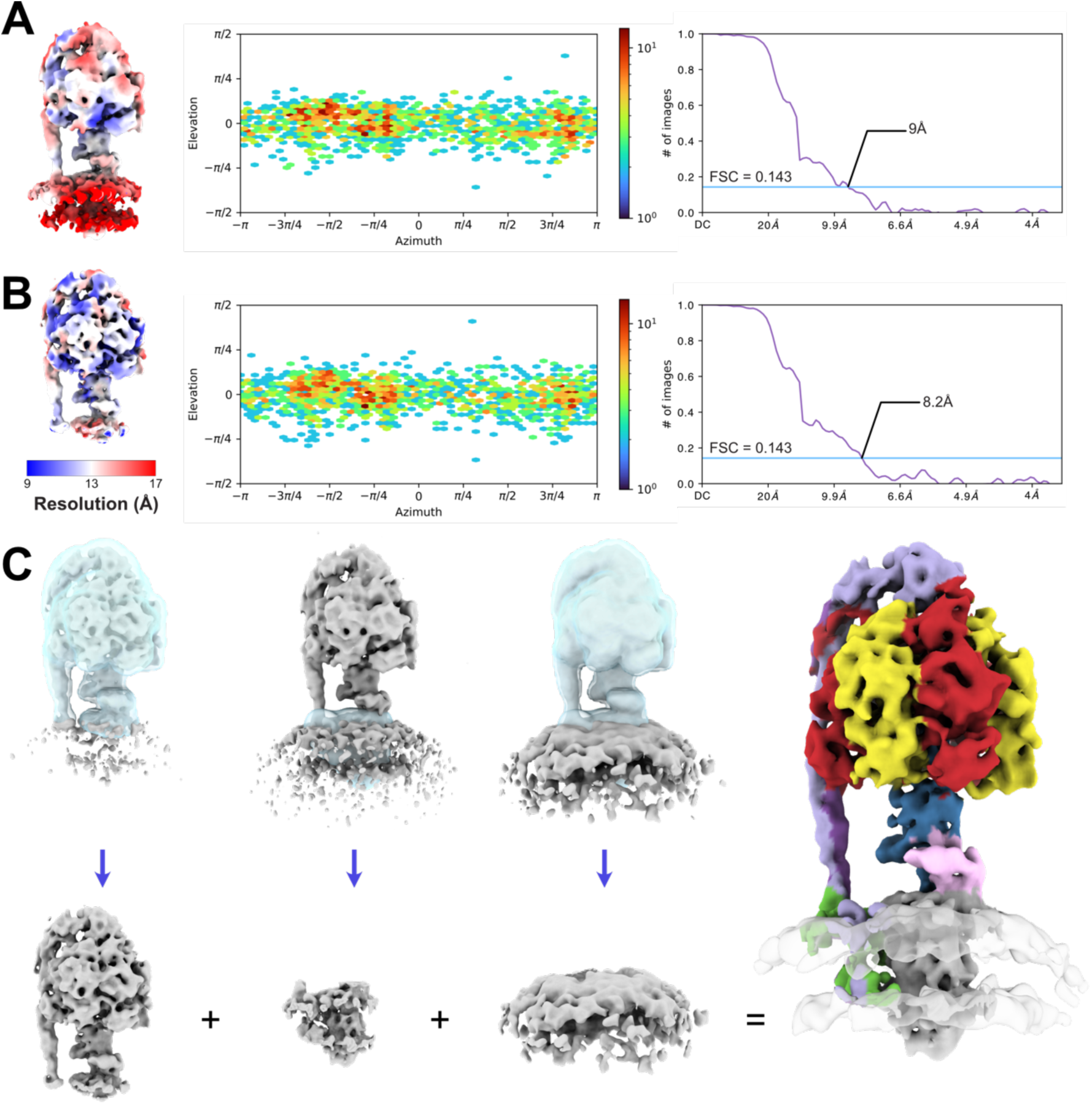
Preparation of a composite map of ATP synthase in native IMVs. **A,** Refinement of an overall map of the ATP synthase showing local resolution (*left*), particle orientation distribution (*middle*), and Fourier shell correlation following a gold standard refinement with correction for the effects of masking (*right*). **B,** Focused refinement of the F_1_ region, including central and peripheral stalks, showing local resolution (*left*), particle orientation distribution (*middle*), and Fourier shell correlation following a gold standard refinement with correction for the effects of masking (*right*). **C,** Combination of the F_1_ region, F_O_ region, and density from the lipid bilayer into a composite map.

**Figure S3.**
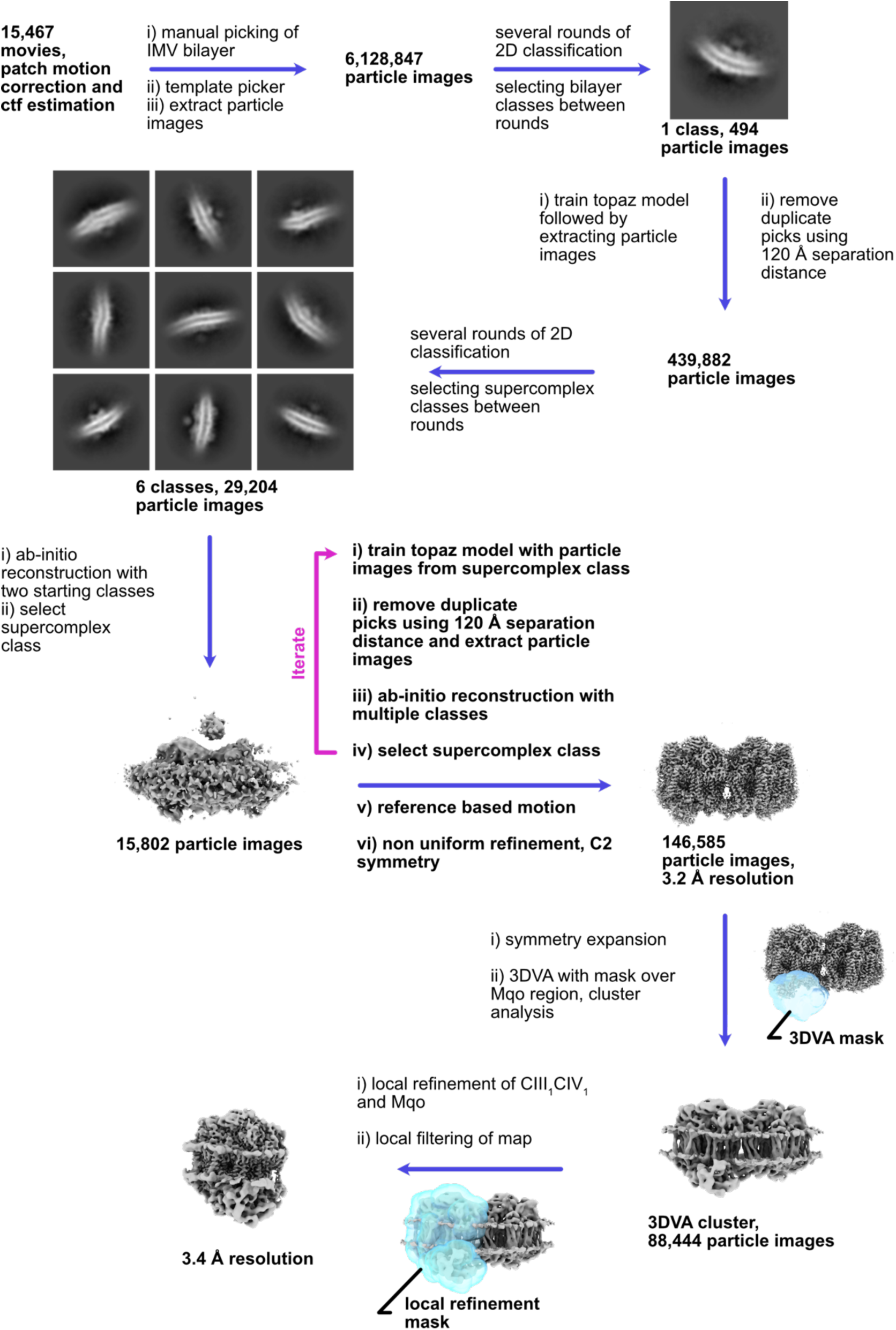
Workflow for calculating map of CIII_2_CIV_2_.

**Figure S4.**
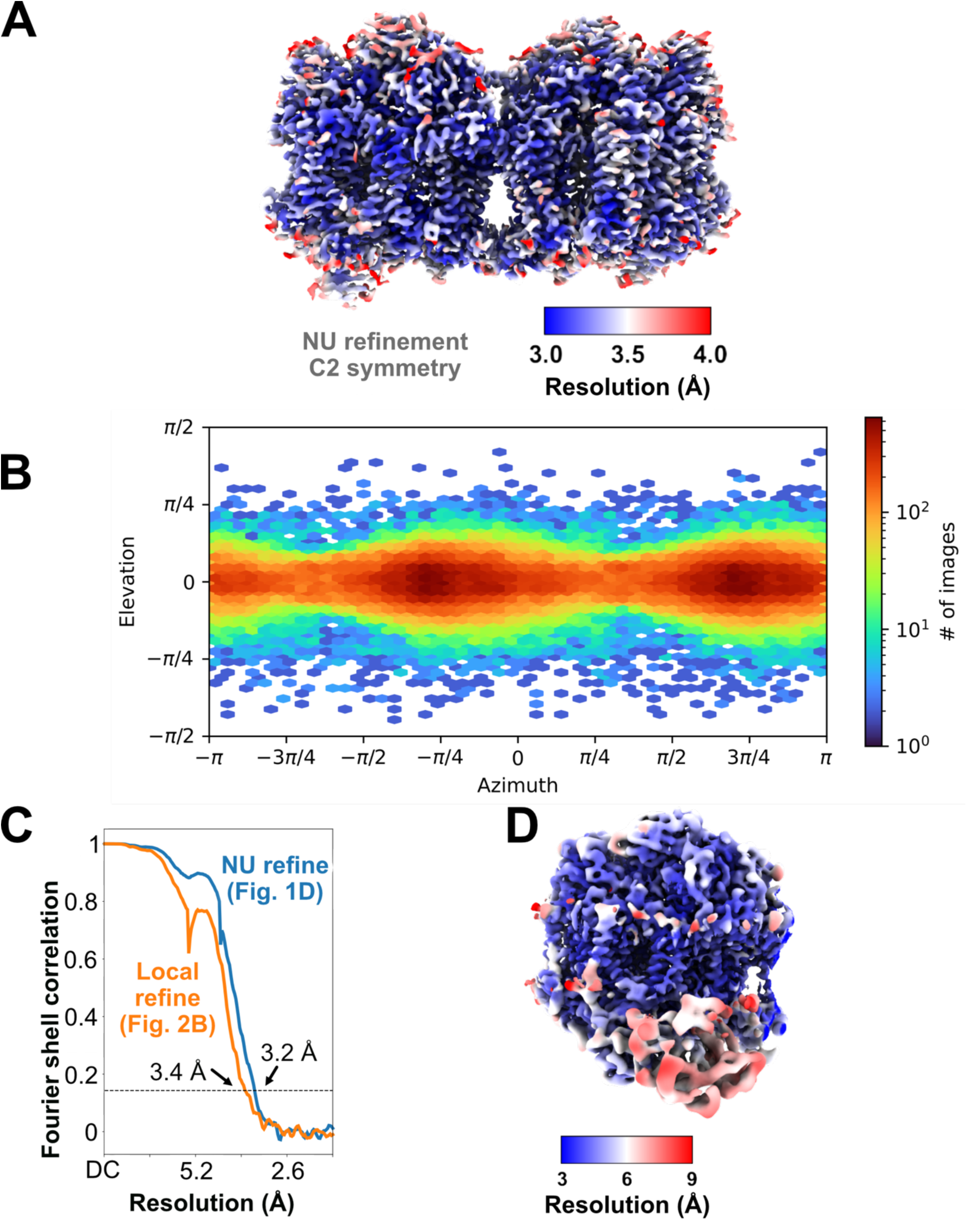
Validation of CIII_2_CIV_2_ map. **A,** Local resolution. **B,** Orientation distribution plot. **C,** Fourier shell correlation following a gold standard refinement with correction for the effects of masking (including focused refinement of a single CIIICIV monomer). **D,** Local resolution for a single CIIICIV monomer.

**Figure S5.**
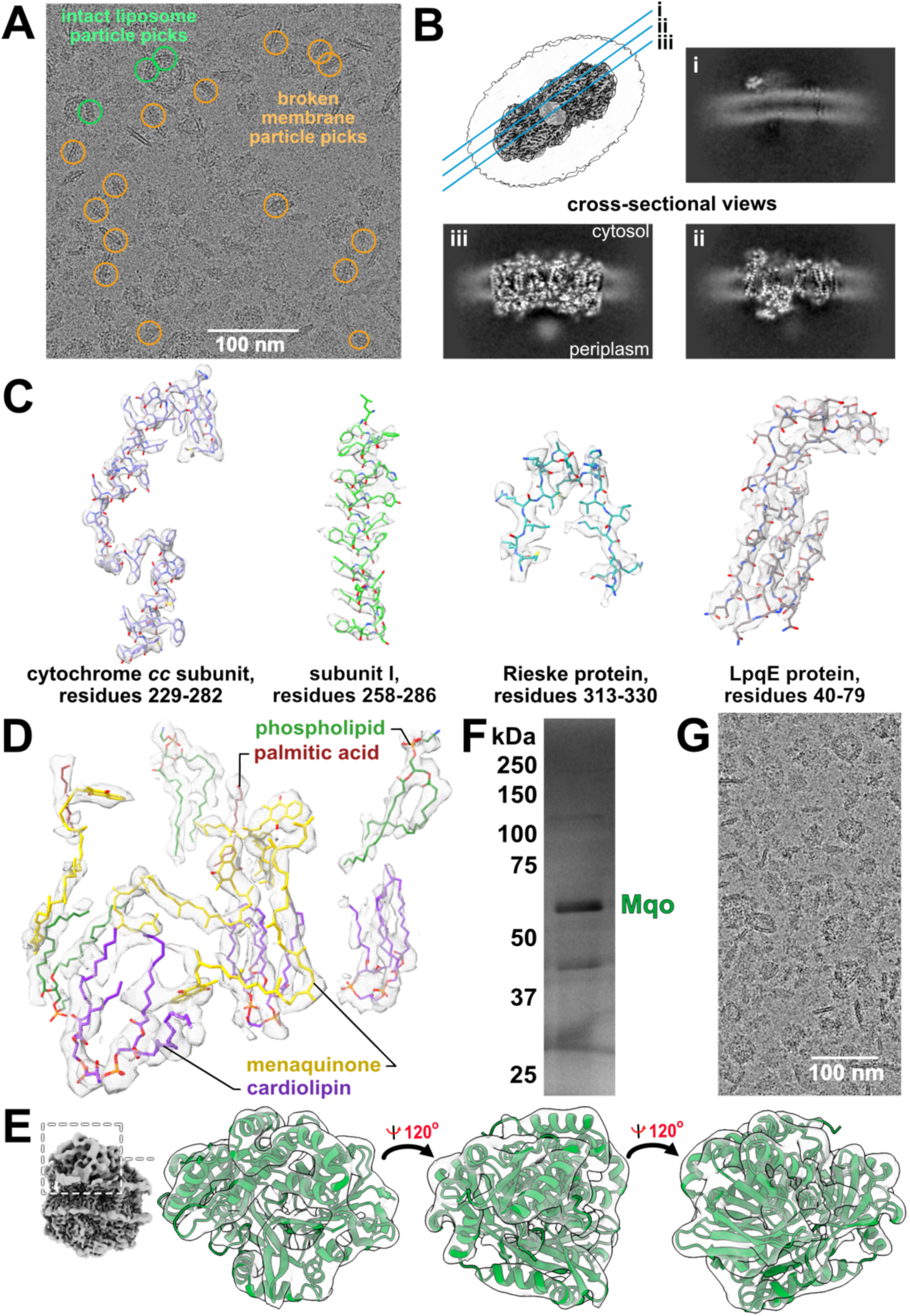
Structure of the Mqo:CIII_2_CIV_2_ complex. **A,** Location of particle images contributing to the 3D map. **B,** Curvature of the lipid bilayer. **C,** Representative high-resolution densities. **D,** Densities for menaquinone and lipids in the map. **E,** Density for Mqo. **F,** Purification of Mqo following detergent extraction. **G,** Isolation of IMVs via 3×FLAG tagged Mqo.

**Supplementary Table 1.** . Cryo-EM data acquisition and image processing.

|  | ATP synthase map | CIII<sub>2</sub>CIV<sub>2</sub> map |
| --- | --- | --- |
| Electron Microscope | Titan Krios |  |
| Camera | Falcon 4i |  |
| Voltage (kV) | 300 |  |
| Nominal Magnification | 75,000 |  |
| Calibrated physical pixel size (Å) | 1.03 |  |
| Total exposure (e/Å<sup>2</sup>) | 40 | 42 |
| Exposure rate (e/pixel/s) | 5.8 | 5 |
| Number of frames | 40 | 29 |
| Defocus range (µm) | 0.7 to 2.4 | 0.9 to 2.5 |
| **Image Processing** |  |  |
| Motion correction software | *MotionCor2* |  |
| CTF estimation software | *cryoSPARC v4.4.0* |  |
| Particle selection software | *cryoSPARC v4.4.0* |  |
| Micrographs used | 8,896 | 15,467 |
| Particle images selected | 3,192 | 146,585 |
| 3D map classification and refinement software | *cryoSPARC v4.4.0* |  |
| EMDB | EMD-47123, EMD-47124 | EMD-46995 |

**Supplementary Table 2.** Atomic model statistics for CIII_2_CIV_2_.

| CIII<sub>2</sub>CIV<sub>2</sub> atomic model statistics |  |
| --- | --- |
| Associated PDB ID | 8D4T |
| PDB ID used for rigid body fit | 7RH5 (Yanofsky et al., 2021) |
| Modelling and refinement software | Chimera1.13.1rc, phenix |
| Protein residues | 6557 |
| Ligand | 9XX:4, 9Y0:6, CDL:16, CU:6, 9YF:8, HEA:4, HEC:4, MQ9:8, HEM:4, PLM:4, FES:2 |
| RMSD bond length (Å) | 0.008 |
| RMSD bond angle (°) | 2.516 |
| Ramachandran outliers (%) | 0.09 |
| Ramachandran favoured (%) | 92.69 |
| Rotamer outliers (%) | 1.15 |
| Clash score | 10.65 |
| MolProbability score | 2.05 |
| EMringer score | 1.43 |

## References

Baugh L, Phan I, Begley DW, Clifton MC, Armour B, Dranow DM, Taylor BM, Muruthi MM, Abendroth J, Fairman JW, Fox D, Dieterich SH, Staker BL, Gardberg AS, Choi R, Hewitt SN, Napuli AJ, Myers J, Barrett LK, Zhang Y, Ferrell M, Mundt E, Thompkins K, Tran N, Lyons-Abbott S, Abramov A, Sekar A, Serbzhinskiy D, Lorimer D, Buchko GW, Stacy R, Stewart LJ, Edwards TE, Van Voorhis WC, Myler PJ. 2015. Increasing the structural coverage of tuberculosis drug targets. Tuberculosis 95:142–148. doi:10.1016/j.tube.2014.12.003

Beites T, O’Brien K, Tiwari D, Engelhart CA, Walters S, Andrews J, Yang HJ, Sutphen ML, Weiner DM, Dayao EK, Zimmerman M, Prideaux B, Desai PV, Masquelin T, Via LE, Dartois V, Boshoff HI, Barry CE, Ehrt S, Schnappinger D. 2019. Plasticity of the Mycobacterium tuberculosis respiratory chain and its impact on tuberculosis drug development. Nat Commun 10:1–12. doi:10.1038/s41467-019-12956-2

Bepler T, Morin A, Rapp M, Brasch J, Shapiro L, Noble AJ, Berger B. 2019. Positive-unlabeled convolutional neural networks for particle picking in cryo-electron micrographs. Nat Methods 16:1153–1160. doi:10.1038/s41592-019-0575-8

Chong SMS, Manimekalai MSS, Sarathy JP, Williams ZC, Harold LK, Cook GM, Dick T, Pethe K, Bates RW, Grüber G. 2020. Antituberculosis Activity of the Antimalaria Cytochrome bcc Oxidase Inhibitor SCR0911. ACS Infect Dis 6:725–737. doi:10.1021/acsinfecdis.9b00408

Coupland CE, Karimi R, Bueler SA, Liang Y, Courbon GM, Di Trani JM, Wong CJ, Saghian R, Youn J-Y, Wang L-Y, Rubinstein JL. 2024. High-resolution electron cryomicroscopy of V-ATPase in native synaptic vesicles. Science 385:168–174. doi:10.1126/science.adp5577

Courbon GM, Palme PR, Mann L, Richter A, Imming P, Rubinstein JL. 2023. Mechanism of mycobacterial ATP synthase inhibition by squaramides and second generation diarylquinolines. EMBO J e113687. doi:10.15252/embj.2023113687

de Carvalho LPS, Fischer SM, Marrero J, Nathan C, Ehrt S, Rhee KY. 2010. Metabolomics of *Mycobacterium tuberculosis* Reveals Compartmentalized Co-Catabolism of Carbon Substrates. Chem Biol 17:1122–1131. doi:10.1016/j.chembiol.2010.08.009

de Souza GA, Arntzen MØ, Fortuin S, Schürch AC, Målen H, McEvoy CRE, van Soolingen D, Thiede B, Warren RM, Wiker HG. 2011. Proteogenomic Analysis of Polymorphisms and Gene Annotation Divergences in Prokaryotes using a Clustered Mass Spectrometry-Friendly Database*. Mol Cell Proteomics 10:M110.002527. doi:10.1074/mcp.M110.002527

Gong H, Gao Y, Zhou X, Xiao Y, Wang W, Tang Y, Zhou S, Zhang Y, Ji W, Yu L, Tian C, Lam SM, Shui G, Guddat LW, Wong LL, Wang Q, Rao Z. 2020. Cryo-EM structure of trimeric Mycobacterium smegmatis succinate dehydrogenase with a membrane-anchor SdhF. Nat Commun 11:1–8. doi:10.1038/s41467-020-18011-9

Gong H, Li Jun, Xu A, Tang Y, Ji W, Gao R, Wang S, Yu L, Tian C, Li Jingwen, Yen H-Y, Man Lam S, Shui G, Yang X, Sun Y, Li X, Jia M, Yang C, Jiang B, Lou Z, Robinson CV, Wong L-L, Guddat LW, Sun F, Wang Q, Rao Z. 2018. An electron transfer path connects subunits of a mycobacterial respiratory supercomplex. Science 362:eaat8923. doi:10.1126/science.aat8923

Guo H, Courbon GM, Bueler SA, Mai J, Liu J, Rubinstein JL. 2021. Structure of mycobacterial ATP synthase with the TB drug bedaquiline. Nature 589:143–147. doi:10.1038/s41586-020-3004-3

Harden SA, Courbon GM, Liang Y, Kim AS, Rubinstein JL. 2024. A simple assay for inhibitors of mycobacterial oxidative phosphorylation. J Biol Chem 300:105483. doi:10.1016/j.jbc.2023.105483

Harold LK, Jinich A, Hards K, Cordeiro A, Keighley LM, Cross A, McNeil MB, Rhee K, Cook GM. 2022. Deciphering functional redundancy and energetics of malate oxidation in mycobacteria. J Biol Chem 298:101859. doi:10.1016/j.jbc.2022.101859

Jumper J, Evans R, Pritzel A, Green T, Figurnov M, Ronneberger O, Tunyasuvunakool K, Bates R, Žídek A, Potapenko A, Bridgland A, Meyer C, Kohl SAA, Ballard AJ, Cowie A, Romera-Paredes B, Nikolov S, Jain R, Adler J, Back T, Petersen S, Reiman D, Clancy E, Zielinski M, Steinegger M, Pacholska M, Berghammer T, Bodenstein S, Silver D, Vinyals O, Senior AW, Kavukcuoglu K, Kohli P, Hassabis D. 2021. Highly accurate protein structure prediction with AlphaFold. Nature 596:583–589. doi:10.1038/s41586-021-03819-2

Kao WC, Kleinschroth T, Nitschke W, Baymann F, Neehaul Y, Hellwig P, Richers S, Vonck J, Bott M, Hunte C. 2016. The obligate respiratory supercomplex from Actinobacteria. Biochim Biophys Acta - Bioenerg 1857:1705–1714. doi:10.1016/j.bbabio.2016.07.009

Kumar R, Sharma P, Chauhan A, Singh N, Prajapati VM, Singh SK. 2024. Malate:quinone oxidoreductase knockout makes Mycobacterium tuberculosis susceptible to stress and affects its in vivo survival. Microbes Infect 26:105215. doi:10.1016/j.micinf.2023.105215

Liang Y, Plourde A, Bueler SA, Liu J, Brzezinski P, Vahidi S, Rubinstein JL. 2023. Structure of mycobacterial respiratory complex I. Proc Natl Acad Sci 120:e2214949120. doi:10.1073/pnas.2214949120

Liang Y, Rubinstein JL. 2023. Structural analysis of mycobacterial electron transport chain complexes by cryoEM. Biochem Soc Trans 51:183–193. doi:10.1042/BST20220611

Målen H, Pathak S, Søfteland T, de Souza GA, Wiker HG. 2010. Definition of novel cell envelope associated proteins in Triton X-114 extracts of Mycobacterium tuberculosis H37Rv. BMC Microbiol 10:132. doi:10.1186/1471-2180-10-132

Marr CR, Benlekbir S, Rubinstein JL. 2014. Fabrication of carbon films with ∼500nm holes for cryo-EM with a direct detector device. J Struct Biol 185:42–47. doi:10.1016/j.jsb.2013.11.002

Mattow J, Schaible UE, Schmidt F, Hagens K, Siejak F, Brestrich G, Haeselbarth G, Müller E-C, Jungblut PR, Kaufmann SHE. 2003. Comparative proteome analysis of culture supernatant proteins from virulent Mycobacterium tuberculosis H37Rv and attenuated M. bovis BCG Copenhagen. ELECTROPHORESIS 24:3405–3420. doi:10.1002/elps.200305601

Mawuenyega KG, Forst CV, Dobos KM, Belisle JT, Chen J, Bradbury EM, Bradbury ARM, Chen X. 2005. Mycobacterium tuberculosis Functional Network Analysis by Global Subcellular Protein Profiling. Mol Biol Cell 16:396–404. doi:10.1091/mbc.e04-04-0329

Molenaar D, Van Der Rest ME, Petrović S. 1998. Biochemical and genetic characterization of the membrane-associated malate dehydrogenase (acceptor) from *Corynebacterium glutamicum*. Eur J Biochem 254:395–403. doi:10.1046/j.1432-1327.1998.2540395.x

Murphy KC, Nelson SJ, Nambi S, Papavinasasundaram K, Baer CE, Sassetti CM. 2018. ORBIT: a New Paradigm for Genetic Engineering of Mycobacterial Chromosomes. mBio 9:e01467–18.

Omini J, Dele-Osibanjo T, Kim H, Zhang J, Obata T. 2024. Is the TCA cycle malate dehydrogenase-citrate synthase metabolon an illusion? Essays Biochem EBC20230084. doi:10.1042/EBC20230084

Punjani A, Fleet DJ. 2021. 3D variability analysis: Resolving continuous flexibility and discrete heterogeneity from single particle cryo-EM. J Struct Biol 213:107702. doi:10.1016/j.jsb.2021.107702

Punjani A, Rubinstein JL, Fleet DJ, Brubaker MA. 2017. cryoSPARC: Algorithms for rapid unsupervised cryo-EM structure determination. Nat Methods 14:290–296. doi:10.1038/nmeth.4169

Punjani A, Zhang H, Fleet DJ. 2020. Non-uniform refinement: adaptive regularization improves single-particle cryo-EM reconstruction. Nat Methods 17:1214–1221. doi:10.1038/s41592-020-00990-8

Safarian S, Opel-Reading HK, Wu D, Mehdipour AR, Hards K, Harold LK, Radloff M, Stewart I, Welsch S, Hummer G, Cook GM, Krause KL, Michel H. 2021. The cryo-EM structure of the bd oxidase from M. tuberculosis reveals a unique structural framework and enables rational drug design to combat TB. Nat Commun 12:5236. doi:10.1038/s41467-021-25537-z

Sarewicz M, Osyczka A. 2015. Electronic connection between the quinone and cytochrome c redox pools and its role in regulation of mitochondrial electron transport and redox signaling. Physiol Rev 95:219–243. doi:10.1152/physrev.00006.2014

Srere PA. 1985. The metabolon. Trends Biochem Sci 10:109–110. doi:10.1016/0968-0004(85)90266-X

Tao X, Zhao C, MacKinnon R. 2023. Membrane protein isolation and structure determination in cell-derived membrane vesicles. Proc Natl Acad Sci 120:e2302325120. doi:10.1073/pnas.2302325120

Vinogradov AD, Kotlyar AB, Burov VI, Belikova YO. 1989. Regulation of succinate dehydrogenase and tautomerization of oxaloacetate. Adv Enzyme Regul 28:271–280. doi:10.1016/0065-2571(89)90076-9

Wang C, Jiang W, Leitz J, Yang K, Esquivies L, Wang X, Shen X, Held RG, Adams DJ, Basta T, Hampton L, Jian R, Jiang L, Stowell MHB, Baumeister W, Guo Q, Brunger AT. 2024. Structure and topography of the synaptic V-ATPase–synaptophysin complex. Nature 631:899–904. doi:10.1038/s41586-024-07610-x

Wang W, Gao Y, Tang Y, Zhou X, Lai Y, Zhou S, Zhang Y, Yang X, Liu F, Guddat LW, Wang Q, Rao Z, Gong H. 2021. Cryo-EM structure of mycobacterial cytochrome bd reveals two oxygen access channels. Nat Commun 12:4621. doi:10.1038/s41467-021-24924-w

Wiseman B, Nitharwal RG, Fedotovskaya O, Schäfer J, Guo H, Kuang Q, Benlekbir S, Sjöstrand D, Ädelroth P, Rubinstein JL, Brzezinski P, Högbom M. 2018. Structure of a functional obligate complex III2IV2 respiratory supercomplex from Mycobacterium smegmatis. Nat Struct Mol Biol 25:1128–1136. doi:10.1038/s41594-018-0160-3

Xiong Y, Chalmers MJ, Gao FP, Cross TA, Marshall AG. 2005. Identification of *Mycobacterium t uberculosis* H37Rv Integral Membrane Proteins by One-Dimensional Gel Electrophoresis and Liquid Chromatography Electrospray Ionization Tandem Mass Spectrometry. J Proteome Res 4:855–861. doi:10.1021/pr0500049

Yanofsky DJ, Di Trani JM, Król S, Abdelaziz R, Bueler SA, Imming P, Brzezinski P, Rubinstein JL. 2021. Structure of mycobacterial CIII2CIV2 respiratory supercomplex bound to the tuberculosis drug candidate telacebec (Q203). eLife 10:e71959. doi:10.7554/eLife.71959

Zheng W, Chai P, Zhu J, Zhang K. 2024. High-resolution In-situ Structures of Mammalian Mitochondrial Respiratory Supercomplexes in Reaction within Native Mitochondria. doi:10.1101/2024.04.02.587796

Zhou S, Wang W, Zhou X, Zhang Y, Lai Y, Tang Y, Xu J, Li D, Lin J, Yang X, Ran T, Chen H, Guddat LW, Wang Q, Gao Y, Rao Z, Gong H. 2021. Structure of Mycobacterium tuberculosis cytochrome bcc in complex with Q203 and TB47, two anti-TB drug candidates. eLife 10:e69418. doi:10.7554/eLife.69418

Zhou X, Gao Y, Wang W, Yang Xiaolin, Yang Xiuna, Liu F, Tang Y, Lam SM, Shui G, Yu L, Tian C, Guddat LW, Wang Q, Rao Z, Gong H. 2021. Architecture of the mycobacterial succinate dehydrogenase with a membrane-embedded rieske FeS cluster. Proc Natl Acad Sci U S A 118:1–6. doi:10.1073/pnas.2022308118

